# *Pseudomonas aeruginosa* induced acute emphysema through bacterial secretion in mice

**DOI:** 10.1101/2020.09.08.287003

**Authors:** Yajie Zhu, Shuming Pan

## Abstract

Pulmonary emphysema is the major pathological feature of chronic obstructive pulmonary disease (COPD). Although the pathogenesis of emphysema is still not completely understood, but until now a bacterial cause has not really been considered. Recently, we found that the secretion from Pseudomonas aeruginosa could cause severe lung emphysema in mice rapidly. Since the bacterium is ubiquitous and secrets proteases, we hypothesized that direct P. aeruginosa airway infection would have a similar effect. To address this issue, we applied a unilateral lung injury model. First, we observed the dynamic pathophysiology change of acute emphysema. P. aeruginosa secretion was extracted and instilled intratracheally into the left lungs of C57BL/6 and C3H/HeJ mice, while the right lungs were saved as self-control. Alveolar diameter and lung compliance were measured. Later, we tested the effect of P. aeruginosa inoculation in normal C57BL/6 mice, immunosuppressed C57BL/6 mice, and C3H/HeJ (TLR4 deficient) mice. P. aeruginosa secretion extract caused acute panacinar emphysema and decreased dynamic lung compliance. Different types of emphysema are transformable. However, the P. aeruginosa infection could only elicit emphysema in immunosuppressed C57BL/6 mice and C3H/HeJ mice, indicating that normal immunity is essential to protect the hosts from emphysema. Emphysema induced by P. aeruginosa in mice recapitulates all the main features of human emphysema and COPD. Our finding filled a major gap in COPD pathogenesis. We believe *P. aeruginosa* is the underlying cause of COPD.

## Introduction

Chronic obstructive pulmonary disease (COPD) is the third leading cause of death globally, characterized by irreversible obstruction of the small airways (bronchiolitis) and destruction of lung parenchyma (emphysema), which lead to air trapping, dyspnea, and even respiratory failure[1]. The pathogenesis of COPD is largely attributed to cigarette smoking induced chronic lung inflammation; however, it could not explain the fact that many COPD patients did not smoke [2], and only 40% of heavy smokers developed emphysema[3]. Current animal models established by passive tobacco smoking or elastase instillation poorly reproduce the main features of human COPD[4]. Accumulating evidence suggests a better understanding of the disease mechanism is needed.

*Pseudomonas aeruginosa* is a gram-negative, aerobic, opportunistic bacterium with a wide distribution. In the lung, *P. aeruginosa* is known to prevalent among immunocompromised hosts, causing cystic fibrosis, bronchiectasis and nosocomial pneumonia [5]. *P. aeruginosa* would colonize in the lung after COPD formation and contribute to the acute exacerbation [6, 7]. It can be isolated from 4-15% of adults with COPD, a number probably greatly underestimated due to this organism’s unique ability to form biofilms, making it difficult to detect[8]. *P. aeruginosa* expresses a variety of virulence factors including endotoxin, exotoxin, elastase, alkaline proteases and hemolysins [9]. The proteases can disrupt epithelial cell tight junctions, cleave collagen, IgG, IgA, complement, and degrade surfactant proteins, causing harm to the lung[10]. However, the manifestation of *P. aeruginosa* pneumonia reported is nonspecific compared with other bacteria[11].

Recently, we discovered accidently that distinguished from Klebsiella pneumoniae, Staphylococcus aureus, Escherichia coli and Proteus mirabilis, the secretion of *P. aeruginosa* could induce severe pulmonary emphysema in murine rapidly. Surprised by this finding, for bacterial factors were never reported to be an initiator in emphysema development[4, 12], we hypothesized that this phenomenon was caused by the proteases secreted by *P. aeruginosa*. Thus, theoretically, *P. aeruginosa* infection shall lead to acute emphysema too. To address the issue, a unilateral lung instillation model of normal C57BL/6, immunosuppressed C57BL/6 and C3H/HeJ mice were used. We observed the dynamic pathology evolution and physiology changes in this study.

## Materials and Methods

### Ethical statement

All experiments referring to the use of animals in this study were approved by the Institutional Animal Care and Use Committee of Shanghai Xinhua Hospital affiliated to Shanghai Jiao Tong University School of Medicine (XHEC-F-2018-047).

### Mice

C57BL/6 mice (10 weeks old) were purchased from Shanghai Sippr-BK laboratory animal Co. Ltd; C3H/HeJ mice (10weeks old) were purchased from Shanghai SLAC laboratory animal Co. Ltd. All mice were accommodated at the Model Animal Research Center of Xinhua Hospital in a specific pathogen-free animal facility under constant temperature and humidity, with sufficient qualified food and water for 1 to 2 weeks before use.

### Isolation and identification of *P. aeruginosa*

*P. aeruginosa* strain was originally separated from sputum, confirmed through mass spectrometry and the strain kept at the bacteriology lab of Xinhua Hospital. To observe the morphology and culture characteristics, the purified strain was streaked on Sheep blood agar plates (Yi Hua, Shanghai) and nutrient agar plates (Comagal, Shanghai) and incubated at 37°C for 24-36 hours.

### Bacterial secretion extraction and identification

1×10^6^ CFU *P. aeruginosa* were streaked on sheep blood agar plate and incubated at 37°C for 24–48 hours. To harvest the *P. aeruginosa* secretion, all colonies on the blood agar plate were scratched down, collected, and dissolved in 5ml PBS then centrifuged at 3000 rcm for 10 minutes. The supernatant was collected, filtered through a 0.22μm PES membrane filter unit (Millex, Merck), and stored at -20□ for later use. The quantification of the supernatant protein concentration was carried through BCA protein assay (Beyotime, China). Protein types in the secretion were identified by protein mass spectrometry. Briefly, the secretion extract went through enzymatic hydrolysis and was analyzed by a Q-Exactive mass spectrometer (Thermo, USA). Proteome Discoverer (v.2.4) was used to search all of the raw data thoroughly against the protein database (UniProt_ *Pseudomonas aeruginosa*). Names and abundances of proteins were acquired as the result.

### Live bacteria preparation

*P. aeruginosa* was streaked on sheep blood agar plate and incubated at 37°C for 8 hours. A proper number of colonies were inoculated in sterile PBS and the turbidity was adjusted to 2 McFarland standards (MCF), which equals 6×10^8^ CFU/ml, by the standard procedure for later use.

### Immunosuppression model

The immunosuppressed model of C57BL/6 mice was established by intraperitoneal administration of dexamethasone sodium phosphate (DEX) 25mg/kg per day for a consecutive 5 days referring to the method described by Rasmussen KR and colleges[13]. Control mice received the same dosage of normal saline intraperitoneally. The immunosuppression status was confirmed on the sixth day by evaluating the organ-to-body weight ratios of the spleen (relative spleen weight)[14]. C3H/HeJ mice were used directly as immunodeficient mice for their lack of Toll-like receptor 4.

### Unilateral lung instillation

In the unilateral lung injury mice model, the left lung was set as the instillation side and the right lung was set as the control side. Mice were anesthetized through injecting 1% pentobarbital sodium (50mg/Kg) intraperitoneally and put to a proper supine position. The anterior cervical hair was removed and the skin disinfected with 75% ethanol. To expose the trachea, a small longitudinal incision was made and endotracheal intubation performed between the second to fourth cricoid cartilage using a 24G intravenous indwelling catheter (Introcan, BRAUN). The 19mm long 24G cannula was then inserted gently into the left main bronchus. A volume of 30ul *P. aeruginosa* secretion extract was delivered into the left lung of each mouse using microliter syringes (Gaoge, Shanghai). Different dosage of live bacteria was inoculated through the same method. After the intratracheal instillation, mice were put back to the mouse cage with a thermal pad underneath, resting in the left recumbent position until full recovery from anesthesia.

### Measurement of lung compliance

Dynamic lung compliances were measured 7 days after unilateral lung instillation of bacterial secretion. The main surgery procedures were the same as above. A 22G intravenous indwelling catheter (Introcan, BRAUN) was used for endotracheal intubation. After exposure of the main trachea, 5mm of the catheter tip was inserted into the main trachea and a ventilator for rodents (Vent Jr, Kent) was connected to the catheter. Target tidal volume range between 0.2-0.5ml was set and the corresponding peak pressure was recorded each (RR=100bpm). Lung dynamic compliance (C_dyn_) was calculated through linear regression by GraphPad Prism 8 software.

### Lung histopathology

Mice were sacrificed at different time points set. The lung was inflated with 4% paraformaldehyde fixative under constant pressure of 15 to 25 cm H_2_O via the trachea and the trachea then tied off. Lung and heart were harvested en bloc and submerged in fixative for approximately 24 hours before the heart was removed and lung was embedded in paraffin blocks for tissue sections (5μm). Hematoxylin and eosin staining were performed subsequently. Slides were scanned (PANNORAMIC, 3DHISTECH), reviewed and evaluated.

### Alveolar mean linear intercept (Lm)

Air space enlargement was measured using the Short & Hennig method, 1967. Briefly, HE stained sections were scanned and reviewed in Caseviewer software. 10 random fields of each right lung and left lung were randomly selected. Equally crossed two lines (both 1000um) were drawn on the scope. The mean linear intercept Lm was then calculated: Lm=N*L/m [15]. In this case, Lm=2*10^4^/m (μm). To better measure how much the left lung air space has enlarged compare to the right lung, Lm Ratio was further calculated, Lm Ratio=Lm (left lung)/ Lm (right lung).

### Statistics

The data were expressed as the mean ± standard error of mean. Statistical analysis was performed in the GraphPad Prism 8 software. Pair-wise comparisons were made using t-tests, whereas multiple comparisons were made using one-way analysis of variance (ANOVA) test. The confidence interval was set at 95% for all tests. Groups of three or four animals were used in each experiment.

## Results

### Identification of *P. aeruginosa*

*P. aeruginosa* was originally isolated and purified from a clinical sputum sample and confirmed through mass spectrometry. The strain exhibited typical *P. aeruginosa* characteristics including strong ginger odor, green metallic sheen, and beta-hemolytic on blood agar with blue-green pigment producing on nutrient agar (**Figure 1**). The *P. aeruginosa* secretion extract was translucid, light green colored with a strong ginger odor. The total protein concentration was 0.35mg/ml.

**Figure 1.**
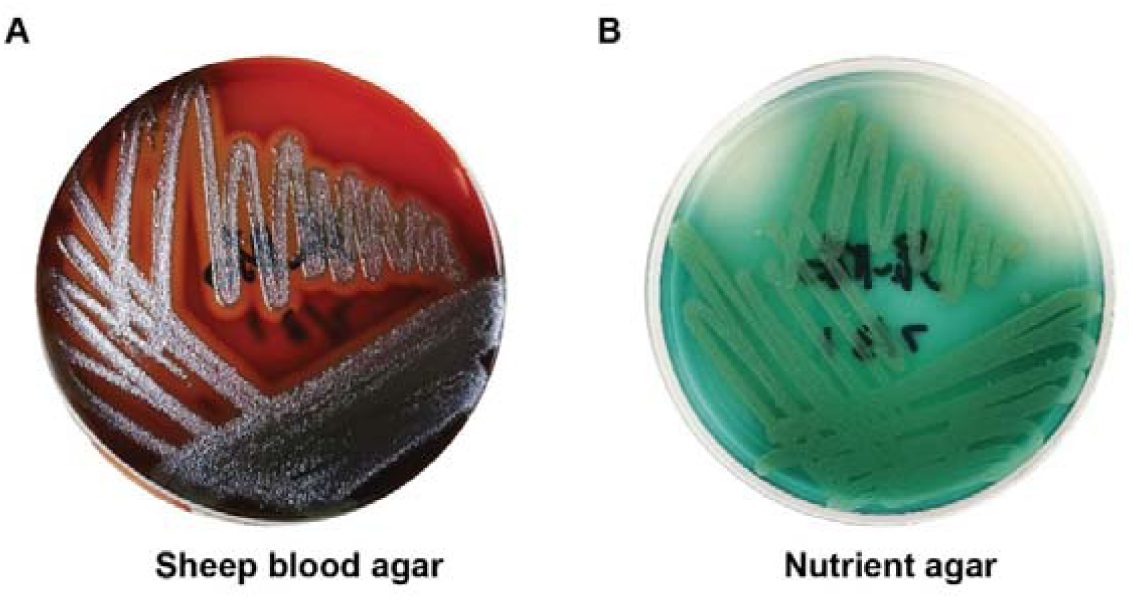
Culture characteristics of the *P. aeruginosa* strain. The strain exhibits typical green metallic sheen, beta-hemolysis, and produces beautiful blue-green pigment.

### *P. aeruginosa* secretion causes different types of emphysema in different mice

Two days after unilateral pulmonary instillation of *P. aeruginosa* secretion, severe parenchymal destruction can be seen, characterized by prominent panacinar emphysema with acute inflammation in the left lung of both C57BL/6 mice and C3H/HeJ mice while the right lung remained mostly unharmed (**Figure 2**). Neutrophil infiltration and erythrocyte diapedesis appeared in both mice group. C3H/HeJ mice had much lighter inflammation response since they are TLR4 deficient and thus blunt in pathogen sensation. Lm Ratio indicates the alveolar of the left lung had enlarged more than three times in C57BL/6 mice, and two times in C3H/HeJ mice (**Figure 3B**).

**Figure 2.**
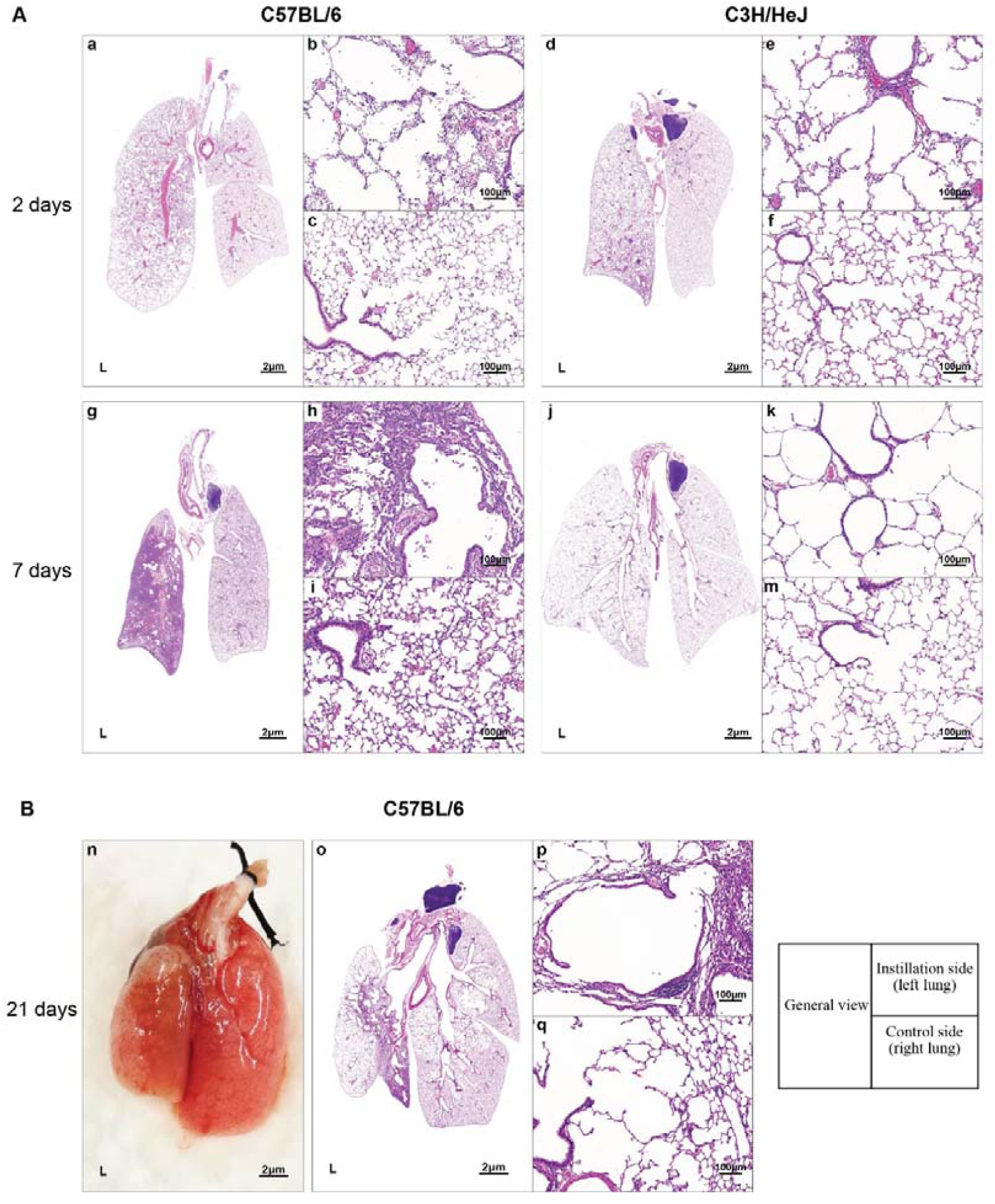
Unilateral pulmonary instillation of *P. aeruginosa* secretion: Two days after instillation, acute panacinar emphysema with inflammation in the left lung of both C57BL/6 mice (b) and C3H/HeJ mice (e). Seven days after instillation, obvious fibrosis and distal emphysema formed in C57BL/6 mice (h) but not in C3H/HeJ (k). Three weeks after stimulation, C57BL/6 mice developed large apical and subpleural bullae (n). Subtypes of emphysema coexisting in the left lung (o, p) and compensatory emphysema developed in the right lung (q). All right lungs served as self-control (c, f, i, m, q); Hematoxylin and eosin (H&E) staining;

**Figure 3:**
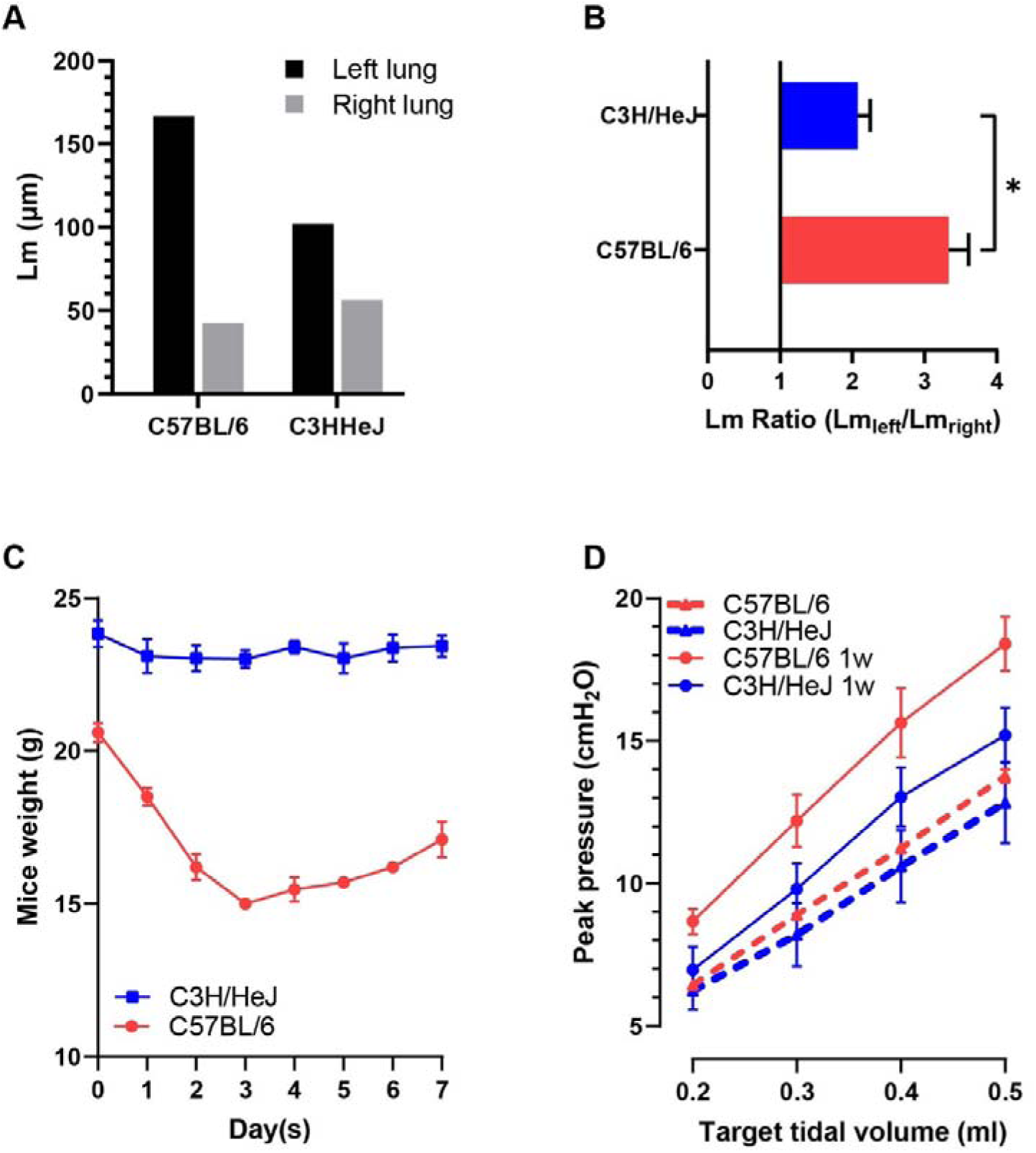
Lm Ratio and dynamic lung compliance: (A) Lm of the C57BL/6 mice (Figure 2-a) and the C3H/HeJ mice (Figure 2-d) 2 days after instillation. (B) Lm ratio indicates the alveolar of the left lung had enlarged more than three times in C57BL/6 group, and two times in C3H/HeJ group. (C) C57BL/6 mice endured obvious weight loss in comparison with C3H/HeJ mice. (D) Dynamic lung compliance of both mice decreased 7 days after modeling. The data represent the mean ± SEM of three animals. *P <.05; **P <.01; ***P<.001;

Seven days after the instillation, inflammation faded and panacinar emphysema was well developed and maintained in C3H/HeJ mice without fibrosis. While interestingly in C57BL/6 mice, the panacinar emphysema had changed to distal emphysema with most of the left lung consolidated. Fibrosis is obvious, inflammation cells can be seen excreting from alveolars and bronchus. To decide if the panacinar emphysema would recur after inflammation resolution, we observed C57BL/6 mice models to three weeks after instillation. Surprisingly, Large apical and subpleural bullae appeared. Histopathology shows all four types of emphysema (centriacinar, panacinar, distal acinar and irregular) coexisting in the left lung and compensatory emphysema developed in the right lung after 21 days (**Figure 2B**).

During the above modeling process, C57BL/6 mice underwent dyspnea, huddle, weight loss and scaphoid abdomen while C3H/HeJ mice showed no obvious symptoms or weight loss (**Figure 3C**).

### *P. aeruginosa* secrets heat-labile proteases

After heated for 80°C, 30mins, the bacterial secretion lost the ability to cause emphysema but still induced neutrophil infiltration (data not shown), indicating different components are responsible for alveolar destruction and inflammation. Protein mass spectrometry confirmed the protein types in the secretion include a high abundance of metal-dependent hydrolase, urocanate hydratase and other enzymes (supplementary material), suggesting the emphysema was caused by heat-labile proteases.

### Dynamic lung compliance decreased after instillation

Seven days after unilateral lung instillation of bacterial secretion, C_dyn_ of C57BL/6 mice decreased from 0.041 to 0.031 ml/cmH_2_O while C_dyn_ of C3H/HeJ mice decreased from 0.045 to 0.036 ml/cmH_2_O. Peak pressure and targeted tidal volume were not well linearly correlated as they used to (**Figure 3D)**.

### *P. aeruginosa* infection induces emphysema in immunocompromised but not immunocompetent mice

Different bacteria dosages, infection time and other conditions in normal C57BL/6 mice were tested, but none of the mice showed noteworthy emphysema (n≥10). A representative case is the mouse was attacked with 15μl of 5 MCF *P. aeruginosa*, which formed a prominent abscess full of *P. aeruginosa* bacteria at the apex of the left lung after 24 hours, but it only caused bronchiectasis-like pathology change in a few areas (**Figure 4)**. Whereas, all C3H/HeJ mice developed homogenous emphysema after live bacteria instillation (n=7). 15μl of 0.5 MCF *P. aeruginosa*—1/10 of the dosage used in C57BL/6 mice—was able to cause obvious centriacinar emphysema in C3H/HeJ. This result reveals that emphysema induced by *P. aeruginosa* is not dosage-dependent.

**Figure 4.**
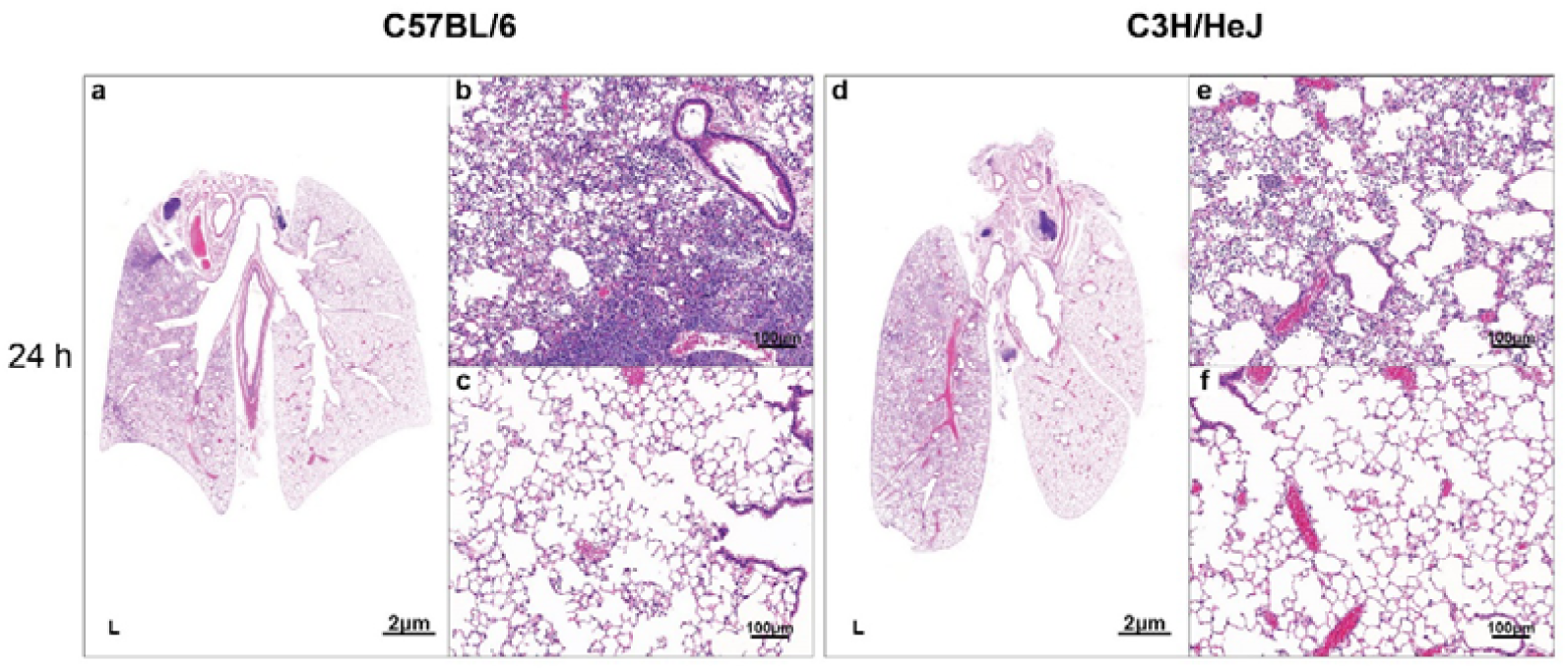
Unilateral live bacteria inoculation: 24 hours after infection, *P. aeruginosa* didn’t induce emphysema in C57BL/6 mice. An abscess full of bacteria at the apex of the left lung formed, but only bronchiectasis-like pathological change (a, b). On the contrary, *P. aeruginosa* inoculation induced prominent centriacinar emphysema in C3H/HeJ mice (d, e). Control lungs (c, f); H&E staining;

In order to decide if the induced emphysema is related to species or immunity differences, we used dexamethasone to suppress the immune reaction of C57BL/6 mice, this immunosuppressed C57BL/6mice, together with C3H/HeJ mice, and normal C57BL/6 mice were infected with 30μl of 2 MCF *P. aeruginosa* for 9 hours (**Figure 5**). Centriacinar pulmonary emphysema occurred in both immunosuppressed C57BL/6 mice and C3H/HeJ mice but not in normal C57BL/6 mice. A comparison of the Lm Ratios between different groups shows that *P. aeruginosa* infection had caused airspace enlargement in two immunocompromised mice groups but alveolar collapse and lung compression in normal C57BL/6 mice group.

**Figure 5.**
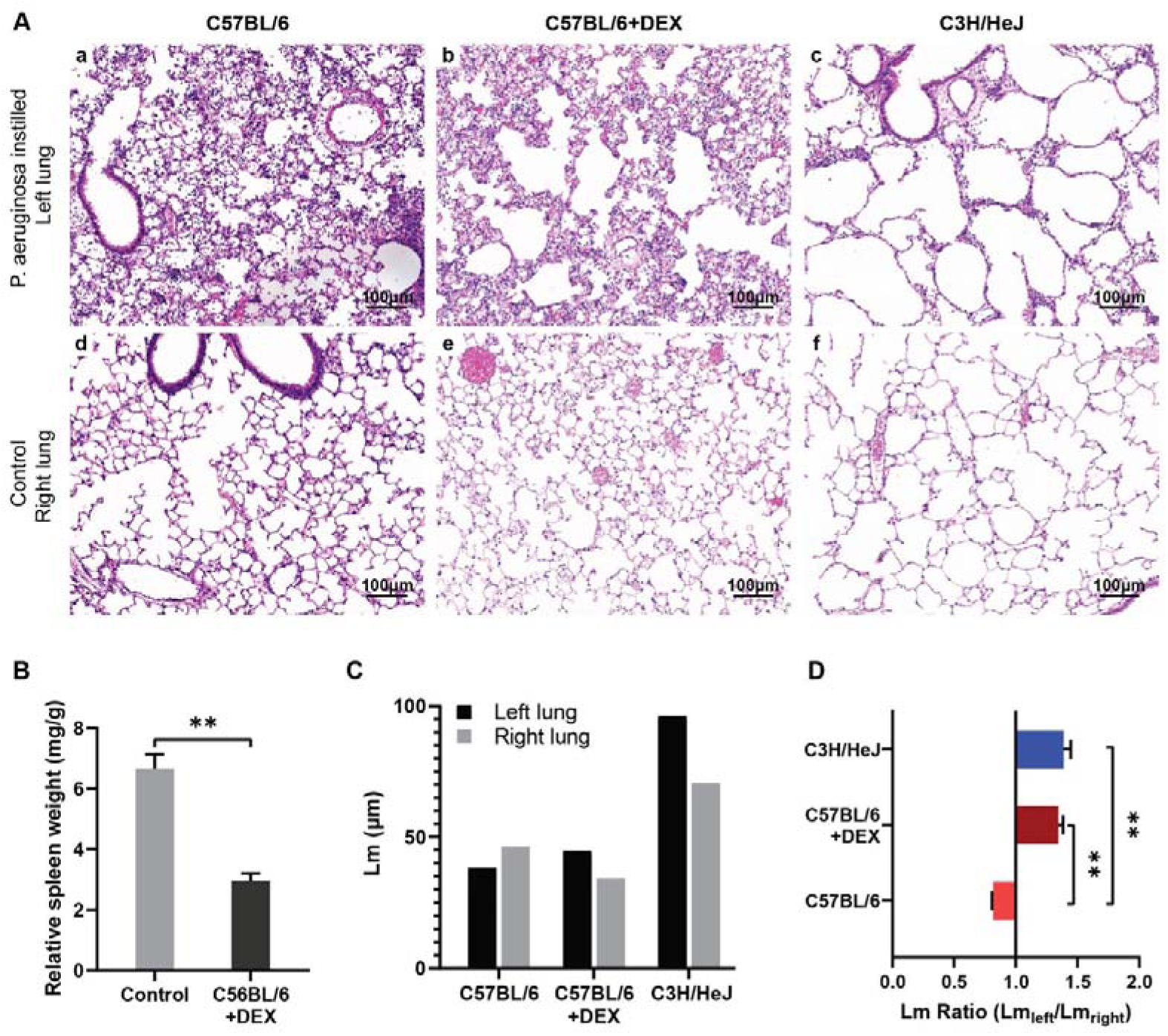
Unilateral live bacteria inoculation in immunosuppressed C57BL/6: (A) Histology of lung of normal C57BL/6 mice, C57BL/6 + DEX mice, and C3H/HeJ mice, Centriacinar pulmonary emphysema developed in both C57BL/6+DEX mice and C3H/HeJ mice, but not in normal C57BL/6 mice. C3HHe/J mice showed the lightest inflammation but most severe emphysema. All mice were inoculated with 30μl of 2 MCF *P. aeruginosa* and sacrificed after 9 hours; H&E staining. (B) The decrease of relative spleen weight of C57BL/6 mice treated with DEX shows successful modeling of hypoimmunity. (C) Lm decreased in C57BL/6 mouse. (D) The Lm ratio between different groups shows *P. aeruginosa* infection caused significant airspace enlargement in two immunocompromised mice groups but airspace compression in the normal C57BL/6 group. The data represent the mean ± SEM of four animals. *P <.05; **P <.01; ***P<.001;

## Discussion

In this study, we applied a unilateral lung injury model and found the secretion of *P. aeruginosa* could cause acute emphysema in mice. Utilizing live *P. aeruginosa* bacteria, we successfully induced emphysema in both immunosuppressed C57BL/6 and C3H/HeJ mice, but not in any C57BL/6 mice with normal immunity. Our findings show that the pathogenesis of COPD can be naturally acute and cigarette smoking irrelevant. Chronic inhalation of noxious particles could have impaired immune function and contributed to COPD formation by creating a circumstance that favors *P. aeruginosa* to survive and thrive [16, 17].

Many researches have pointed out P. aeruginosa is closely related to COPD[18, 19]. *P. aeruginosa* is a nonfastidious bacterium that can proliferate in distilled water and is one of the strongest antibiotic-resistant bacteria[10]. The secretion of *P. aeruginosa* is a mixture of metal-dependent hydrolase, enzymes, and bacterial endo/exotoxins which elicits strong inflammation in normal mice. Interestingly, it seems the neutrophils played a positive role in preventing emphysema formation. Immunosuppressed C57BL/6 as well as TLR4 deficient C3H/HeJ mice who had lighter inflammatory response and neutrophils infiltration than C57BL/6 mice, exhibited more severe airspace dilation.

Protease–antiprotease imbalance as a result of cigarette smoking induced chronic inflammation is the potential mechanism of COPD, based on the observation that patients with a genetic deficiency of α1-anti-trypsin have a predisposition to develop emphysema[3]. Emphysema caused by lung instillation of porcine pancreatic elastase as a widely used disease model supported this theory[20]. Many studies found elevated matrix metalloproteinases (MMPs) concentration and increased elastin degradation in COPD patients [21-23]. Neutrophil or macrophage elastases were held culprit[24]. But it seems that the proteolytic enzymes secreted by various bacteria had not been taken into account. Proteases secretion is a common feature in *P. aeruginosa* and other bacteria[25], however, it didn’t occur to us before that the excessive elastase may come from an outer pathogen rather than our own neutrophils.

Whereas, most researches conducted used animals with normal immunity. α1-anti-trypsin secreted by local macrophages and liver is a major inhibitor of proteases normally presented in serum and tissue fluids. If the immunity is unimpaired, proper inflammation response could increase α1-anti-trypsin level to eliminate the destructive effects caused by bacteria proteases[12]. Our research supported this concept by demonstrating that P. aeruginosa infection only caused limited bronchiectasis in normal mice. But competent immunity does not guarantee a waiver from emphysema. Excessive proteases stimulation proved effective in all mice regardless of background[26]. It is reasonable to conclude that emphysema formation is mainly determined by bacterial proteases dosage and secondly influenced by immunity status.

Centriacinar and panacinar emphysema are most commonly found in COPD patients and regarded as distinct entities. Centriacinar emphysema is closely associated with tobacco smoking while panacinar emphysema is usually associated with α1-anti-trypsin deficiency[27, 28]. However, we found the subtypes of emphysema are related and transformable to each other. In C3H/HeJ mice, the acute emphysema transited to stable irreversible panacinar emphysema while in normal C57BL/6 mice, it went through dynamic change interweaved with inflammation and developed to other subtypes of emphysema as well as bullae. At the same time, fibrosis in the lung has nothing to do with emphysema but is determined by the immune reaction of the hosts. If the stimulus is not persistent, fibrosis would gradually disperse and emphysema would recur.

The difference of air space volume is hard to detect since lung tissue has a nature of instability. In our study, even lungs were carefully inflated with fixative under constant pressure, the many steps in the histopathology preparation process still impacted the final pressure, which can be seen from the variation of alveolar diameters of control lung between different mouse. We solved this difficulty by utilizing the unilateral instillation model which were readily used in a few studies[29, 30]. A mouse model of unilateral injury can provide valuable self-control, eliminate systemic influence, and reduce the use of lab animals. By ensuring the pressure is equal within the lung, minor changes could be detected and compared. The Lm Ratio of left to right lung further revealed clearly how much the airspace has enlarged or compressed.

However, a unilateral lung injury model is not perfect. Sometimes contamination of the control lung caused by the sneeze or cough of mice is not totally avoidable, especially in bacterial infection models. Thus, the histology changes in the main bronchus of the control lung should be interpreted carefully. We tried to do more quantitative assessment of the lung structure according to standard recommendation[31], but found it is impractical when a heavy inflammation occurs (Figure 2h). Due to this reason, we had to sacrifice bacteria inoculated mice after 9 hours, before the inflammatory response makes it impossible to measure Lm.

On the contrary to the current paradigm taking lung infection as no participant in emphysema initiation but only contribute to exacerbation, we discovered for the first time that *P. aeruginosa* caused acute emphysema through bacterial proteases in immunodeficient animals, which is in comply with its predilection to infect immunocompromised patients. Emphysema induced by P. aeruginosa in mice recapitulates all the main features of human emphysema and COPD. We believe *P. aeruginosa* is the underlying cause of COPD. In the future, strategies aimed at developing a vaccine or specialized antibiotics targeting *P. aeruginosa* may be useful for COPD prevention and treatment.

## Acknowledgements

We would like to thank Dr. Feng Chen and Dr. Jingxian Liu (Bacteriology Lab, Xinhua Hospital) as well as Professor Yong Zhang (Department of Immunology, SJTUSM) and Professor Min Wang (Department of Histology and Embryology, SJTUSM) for their skilled advice and assistance;

