## supplementary material for "*Pseudomonas aeruginosa* induced acute emphysema through bacterial secretion in mice"

Protein Mass Spectrometry result of Pseudomonas aeruginosa secretion

| Accession | Description | MW [kDa] | Abundance: F1: Sample |
|---|---|---|---|
| A0A367M2A4 | Metal-dependent hydrolase | 19.2 | 132653.0313 |
| A0A6B0IXI3 | Heme iron utilization protein | 27 | 1165450.375 |
| A0A485CDN9 | Urocanate hydratase | 35.6 | 3348934 |
| A0A485ECM7 | Fumarylacetoacetase | 42.1 | 6335664.5 |
| A0A6H1QMR3 | Inorganic diphosphatase | 19.4 | 1638442.75 |
| A0A5E8W4V4 | TonB-dependent copper receptor | 79.3 | 841702.875 |
| A0A0A8RL96 | Metalloproteinase outer membrane protein | 54.3 | 2403733.75 |
| A0A5K1SF60 | Poly(3-hydroxyalkanoate) granule-associated protein PhaF (Fragment) | 24.7 | 256225.875 |
| A0A1C7BPK6 | Uncharacterized protein | 51.2 | 839050.4375 |
| V6ADM5 | Type I restriction enzyme R Protein | 112.6 | |
| A0A2S5IEA5 | NCS2 family permease | 47 | 301845.125 |
| A0A485GKT5 | Polyhydroxyalkanoate synthesis protein PhaF | 31.1 | 185908.1094 |
| A0A3S3TAW3 | Transposase | 72.6 | 130542.4063 |
| A0A6H1QRX8 | Transcriptional regulator | 36.1 | 217602.3438 |
